# Reducing susceptibility distortion related image blurring in diffusion MRI EPI data

**DOI:** 10.1101/2021.06.07.447406

**Authors:** Ian A. Clark, Martina F. Callaghan, Nikolaus Weiskopf, Eleanor A. Maguire, Siawoosh Mohammadi

**Author notes:** **Correspondence:** Eleanor Maguire, Wellcome Centre for Human Neuroimaging, UCL Queen Square Institute of Neurology, University College London, 12 Queen Square, London, WC1N 3AR, UK;, Siawoosh Mohammadi, Institute of Systems Neuroscience, University Medical Centre Hamburg-Eppendorf, Hamburg, Germany.

## Abstract

Diffusion magnetic resonance imaging (MRI) is an increasingly popular technique in basic and clinical neuroscience. One promising application is to combine diffusion MRI with myelin maps from complementary MRI techniques such as multi-parameter mapping (MPM) to produce g-ratio maps that represent the relative myelination of axons and predict their conduction velocity. Statistical Parametric Mapping (SPM) can process both diffusion data and MPMs, making SPM the only widely accessible software that contains all the processing steps required to perform group analyses of g-ratio data in a common space. However, limitations have been identified in its method for reducing susceptibility-related distortion in diffusion data. More generally, susceptibility-related image distortion is often corrected by combining reverse phase-encoded images (blip-up and blip-down) using the arithmetic mean (AM), however, this can lead to blurred images. In this study we sought to (1) improve the susceptibility-related distortion correction for diffusion MRI data in SPM; (2) deploy an alternative approach to the AM to reduce image blurring in diffusion MRI data when combining blip-up and blip-down EPI data after susceptibility-related distortion correction; and (3) assess the benefits of these changes for g-ratio mapping. We found that the new processing pipeline, called consecutive Hyperelastic Susceptibility Artefact Correction (HySCO) improved distortion correction when compared to the standard approach in the ACID toolbox for SPM. Moreover, using a weighted average (WA) method to combine the distortion corrected data from each phase-encoding polarity achieved greater overlap of diffusion and more anatomically faithful structural white matter probability maps derived from minimally distorted multi-parameter maps as compared to the AM. Third, we showed that the consecutive HySCO WA performed better than the AM method when combined with multi-parameter maps to perform g-ratio mapping. These improvements mean that researchers can conveniently access a wide range of diffusion-related analysis methods within one framework because they are now available within the open-source ACID toolbox as part of SPM, which can be easily combined with other SPM toolboxes, such as the hMRI toolbox, to facilitate computation of myelin biomarkers that are necessary for g-ratio mapping.

## INTRODUCTION

Diffusion magnetic resonance imaging (MRI) is an increasingly popular method in neuroscience and clinical research. Diffusion MRI permits visualisation of the mobility of water molecules within brain tissue, and provides measures of microstructural integrity and anatomical connectivity (Conturo et al., 1999; Shimony et al., 1999; Le Bihan et al., 2001; Kaden et al., 2007). Knowledge of how different brain regions communicate is essential for understanding brain anatomy and its relationship with cognitive processes (Jbabdi et al., 2015; Hodgetts et al., 2017; Assaf et al., 2019; Movahedian Attar et al., 2020), and diffusion imaging in the clinic can be used for various purposes including the characterisation of strokes and tumours (Neumann-Haefelin et al., 1999; Bergui et al., 2001).

However, diffusion MRI data are generally acquired with echo planar imaging (EPI; Turner and Le Bihan, 1990) – a fast MRI acquisition technique that is prone to spatial distortions related to off-resonance effects (Jezzard and Balaban, 1995). It is, therefore, important to correct spatial distortions in diffusion data to provide as accurate a representation as possible of the true anatomy. Fortunately, various methods to address spatial distortions in diffusion MRI are available. Field inhomogeneities due to magnetic susceptibility variations can be measured with additional MRI sequences for field mapping (Jezzard and Balaban, 1995; Reber et al., 1998; Hutton et al., 2002) or estimated from the distorted images themselves using physical models (Chang and Fitzpatrick, 1992). A popular and efficient approach uses two EPI images with reversed phase-encoding directions – also known as “blip-up” and “blip-down” images (Figure 1A) – to estimate the field map and correct the susceptibility-related distortions (Andersson et al., 2003; Weiskopf et al., 2005; Holland et al., 2010; Ruthotto et al., 2012; Breman et al., 2020; other susceptibly distortion correction methods are also available, e.g. Irfanoglu et al., 2015; Schilling et al., 2020). The issue remains, however, that even after distortion correction is applied, the corrected diffusion MRI images can suffer from additional image blurring artefacts because the compression of voxels leads to a loss of information. As illustrated in Figure 1, blip-up and blip-down images are often combined by calculating the arithmetic mean (AM) of the two images to provide a single diffusion image (Figure 1B, top).

**FIGURE 1.**
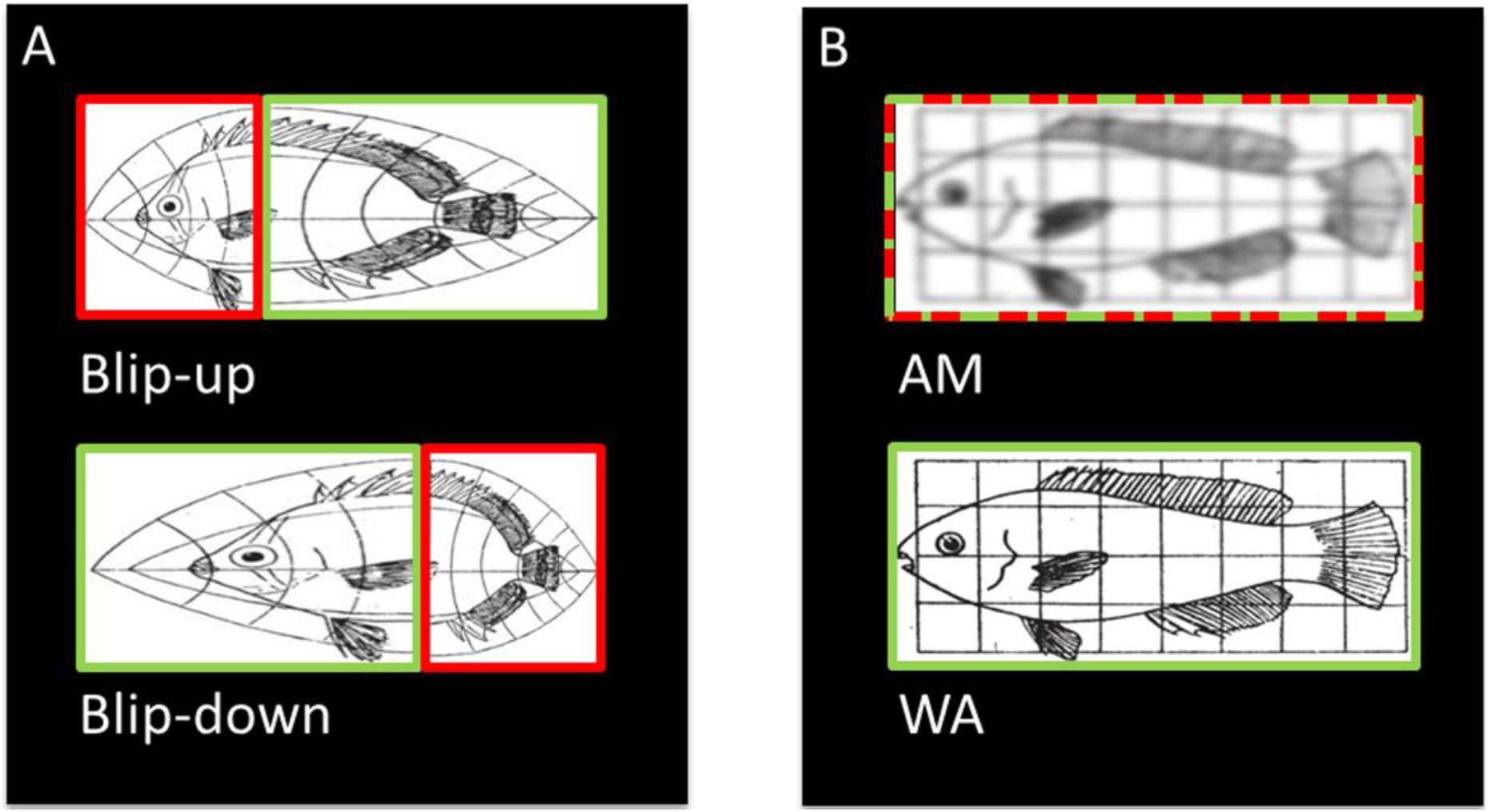
Illustration (modified from Jones and Mahadevan, 2013) of the origin of blurring artefacts after combining distortion corrected diffusion data, and how blurring can be avoided. **(A)** Susceptibility-related distortions in two EPI images acquired with reverse phase-encoding directions (i.e. “blip-up” and blip-down” images) can lead to localized squeezing and stretching. Importantly, where one image is squeezed (red box, top) the reverse phase-encoded image (green box, bottom) is stretched. **(B)** After distortion correction and resampling, the two images are typically combined into an arithmetic mean (AM, red-green-dashed box, top). Here, we propose an alternative combination, the weighted average (WA, green box, bottom), where only the stretched boxes are combined, which retains the spatial information. Therefore, spatial information is lost in the AM combination leading to blurring artefacts whereas it is retained in the WA combination.

Avoiding residual spatial distortions and image blurring is particularly important when combining diffusion MRI with other quantitative MRI contrasts that can reveal additional information that would otherwise not be available (Mohammadi and Callaghan, 2018). For example, multi-parameter mapping (MPM; Weiskopf et al., 2013) is an emerging neuroimaging technique that provides a comprehensive approach to acquiring multiple quantitative biomarkers, namely, the longitudinal relaxation rate *R*_1_, proton density *PD*, effective transverse relaxation rate 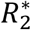, and magnetization transfer saturation *MT*_sat_ (Tabelow et al., 2019). These quantitative parameters contain specific and complementary information regarding the microstructural properties of brain tissue, such as the extent of myelination, iron and water concentration (e.g. Weiskopf et al., 2015; Kirilina et al., 2020). Combining diffusion MRI with MPM thus allows for the examination of microstructural information related to the neuronal conduction velocity in the form of the axonal g-ratio (Mohammadi et al., 2015a; Mohammadi and Callaghan, 2020). Furthermore, by combining diffusion MRI and MPM structural maps, it becomes possible to perform multivariate analyses with biomarkers sensitive to complementary tissue features (Draganski et al., 2011; Chowdhury et al., 2013). While various methods using reverse phase-encoding are available to efficiently reduce susceptibility-related distortions (for example, see Gu and Eklund, 2019), these implementations often focus only on diffusion MRI data. As such, to combine diffusion MRI with other quantitative MRI data, such as MPM for g-ratio mapping, the use of multiple toolboxes implemented in different software packages, is currently required.

FSL, together with TBSS for group analyses, is probably the most widely used package for diffusion MRI (Smith et al., 2004), with TOPUP for distortion correction (Andersson et al., 2003) and eddy for motion and eddy current correction (Andersson and Sotiropoulos, 2016). To examine the g-ratio, however, the diffusion data have to be combined with a quantitative MRI biomarker for the myelin compartment, requiring laboratory-specific in-house software that is not widely available (Stikov et al., 2015; Cercignani et al., 2017; Mancini et al., 2018). In another analysis package, SPM (www.fil.ion.ucl.ac.uk/spm/), the ACID toolbox (www.diffusiontools.com) can be used for distortion correction using Hyperelastic Susceptibility Artefact Correction (HySCO2; Ruthotto et al., 2012; Ruthotto et al., 2013; Macdonald and Ruthotto, 2018) and motion and eddy current correction using ECMOCO (Mohammadi et al., 2010; Mohammadi et al., 2015b). Moreover, the hMRI toolbox, also in SPM (www.hmri.info; Tabelow et al., 2019), provides an easy-to-use framework to generate quantitative structural maps from MPM data. SPM, therefore, has the advantage of being able to process both diffusion and MPM data, making SPM the only widely accessible software that contains all the processing steps required to perform group analyses of g-ratio data in a common space. However, limitations have been identified in the ability of the HySCO2 methodology to reduce susceptibility-related distortion (Gu and Eklund, 2019), highlighting a need for improvement. Moreover, the HySCO2 method uses the AM to combine the oppositely phase-encoded data after correction, meaning that blurring artefacts are typically present.

Here, we sought to improve the pipeline for the HySCO2-based distortion correction, an approach we call consecutive HySCO, and to employ a different method to combine the corrected blip-up and blip-down images to reduce image blurring by using a weighted average (WA). We achieved this by down-weighting regions that were squeezed (e.g. in the blip-up images) while up-weighting the same (stretched) regions in the opposite (blip-down) image. To determine regions that were stretched and squeezed, we used the Jacobian of the HySCO2 estimated field map. We compared the performance of the proposed WA approach to the standard AM combination used by HySCO2 in SPM. Where appropriate, we also benchmarked these changes against the widely available default pipeline in FSL (as used in the human connectome project; Glasser et al., 2013). Finally, given that the overall goal was to have all the processing steps necessary to perform g-ratio analysis within one (SPM) package, we also assessed whether there was any benefit of using the WA corrected diffusion data over the AM corrected data for g-ratio mapping.

## MATERIALS AND METHODS

### Participants

Eighteen participants took part in the study. They were aged between 20 and 40 years old, reported no psychological, psychiatric, neurological or behavioural health conditions. The mean age of the sample was 30.0 years (SD = 6.07) and included 9 females and 9 males. Participants were reimbursed £10 per hour for taking part which was paid at study completion. All participants gave written informed consent and the study was approved by the University College London Research Ethics Committee.

### Diffusion MRI data acquisition

Diffusion-weighted images were collected on Siemens Magnetom 3T TIM Trio systems using 32 channel head coils. The protocol used was the Multi-Band Accelerated EPI Pulse Sequence developed by the Centre for Magnetic Resonance Research at the University of Minnesota (R012a-c, R013a on VB17, https://www.cmrr.umn.edu/multiband/; Feinberg et al., 2010; Xu et al., 2013). Acquisition parameters were: resolution = 1.7 mm isotropic; FOV = 220 mm × 220 mm × 138 mm; 60 directions with 6 interleaved b0 images for a total of 12 b0 images in each of the blip-up and blip-down phase encoding directions, echo time (TE) = 112 ms, repetition time (TR) = 4.84s, with a multiband acceleration factor of 3. The sequence was performed 4 times – twice with b-values of 1000 s/mm^2^ and twice with b-values of 2500 s/mm^2^. The first acquisition of each set of b-values was performed with phase-encoding in the anterior to posterior direction (blip-up), the second in the posterior to anterior direction (blip-down). The total acquisition time was 22 minutes.

### Structural MRI data acquisition

Whole brain structural maps of magnetisation transfer (MT) saturation, at an isotropic resolution of 800μm, were derived from an MPM quantitative imaging protocol (Weiskopf et al., 2013; Callaghan et al., 2015; Callaghan et al., 2019). This protocol consisted of the acquisition of three multi-echo gradient-echo acquisitions with either PD, T1 or MT weighting. Each acquisition had a TR of 25 ms. PD weighting was achieved with an excitation flip angle of 6°, which was increased to 21° to achieve T1 weighting. MT weighting was achieved through the application of a Gaussian RF pulse 2 kHz off resonance with 4ms duration and a nominal flip angle of 220°. This acquisition had an excitation flip angle of 6°. The field of view was 256mm head-foot, 224mm AP, and 179mm right-left. The multiple gradient echoes per contrast were acquired with alternating readout gradient polarity at eight equidistant echo times ranging from 2.34 to 18.44ms in steps of 2.30ms using a readout bandwidth of 488 Hz/pixel. Only six echoes were acquired for the MT weighted volume to facilitate the off-resonance pre-saturation pulse within the TR. To accelerate the data acquisition, partially parallel imaging using the GRAPPA algorithm was employed in each phase-encoded direction (anterior-posterior and right-left) with forty integrated reference lines and a speed up factor of two. Calibration data were also acquired at the outset of each session to correct for inhomogeneities in the RF transmit field (Lutti et al., 2010; Lutti et al., 2012). The total acquisition time was 27 minutes.

### Diffusion MRI distortion correction pipelines

#### Standard HySCO pipeline

The diffusion MRI data were first processed using the standard pre-processing pipeline available in the ACID toolbox (www.diffusiontools.com) within SPM12 (www.fil.ion.ucl.ac.uk/spm). Each diffusion image (regardless of b-value or blip-up / blip-down acquisition) was separately corrected first for motion and eddy current artefacts (Mohammadi et al., 2010), and then for susceptibility-related distortion artefacts using the HySCO2 module (Ruthotto et al., 2012; Ruthotto et al., 2013; Macdonald and Ruthotto, 2018). HySCO is a tool for the hyperelastic susceptibility correction of diffusion data that corrects geometric deformations and intensity modulations in EPI images caused by susceptibility-related field-inhomogeneities. To this end, HySCO takes a tailored variational image registration problem and incorporates a physical distortion model that is aimed at minimising the distance of two oppositely distorted images subject to invertibility constraints.

#### The new consecutive HySCO pipeline

As with the standard HySCO pipeline, each diffusion image (regardless of b-value or blip-up / blip-down acquisition) was first separately corrected for motion and eddy current artefacts and then for susceptibility-related distortion artefacts using the HySCO2 module. Next, the “mean_b0” of the blip-up data was co-registered to the “mean_b0” of the blip-down dataset using a rigid-body transformation (spm_coreg) and the estimated transformation was applied to the full the blip-down dataset. Tensor fitting using all the b-values (Mohammadi et al., 2013) was then performed separately on each of the distortion corrected blip-up and blip-down datasets to estimate Fractional Anisotropy (FA) maps. These FA images were then brain-masked using ACID. Finally, HySCO2 was repeated but now using the distortion corrected and brain-masked FA maps as input instead of b0 images; the second HySCO2 field map being consecutively applied to the “pre-corrected” diffusion MRI data (Figure 2).

**FIGURE 2.**
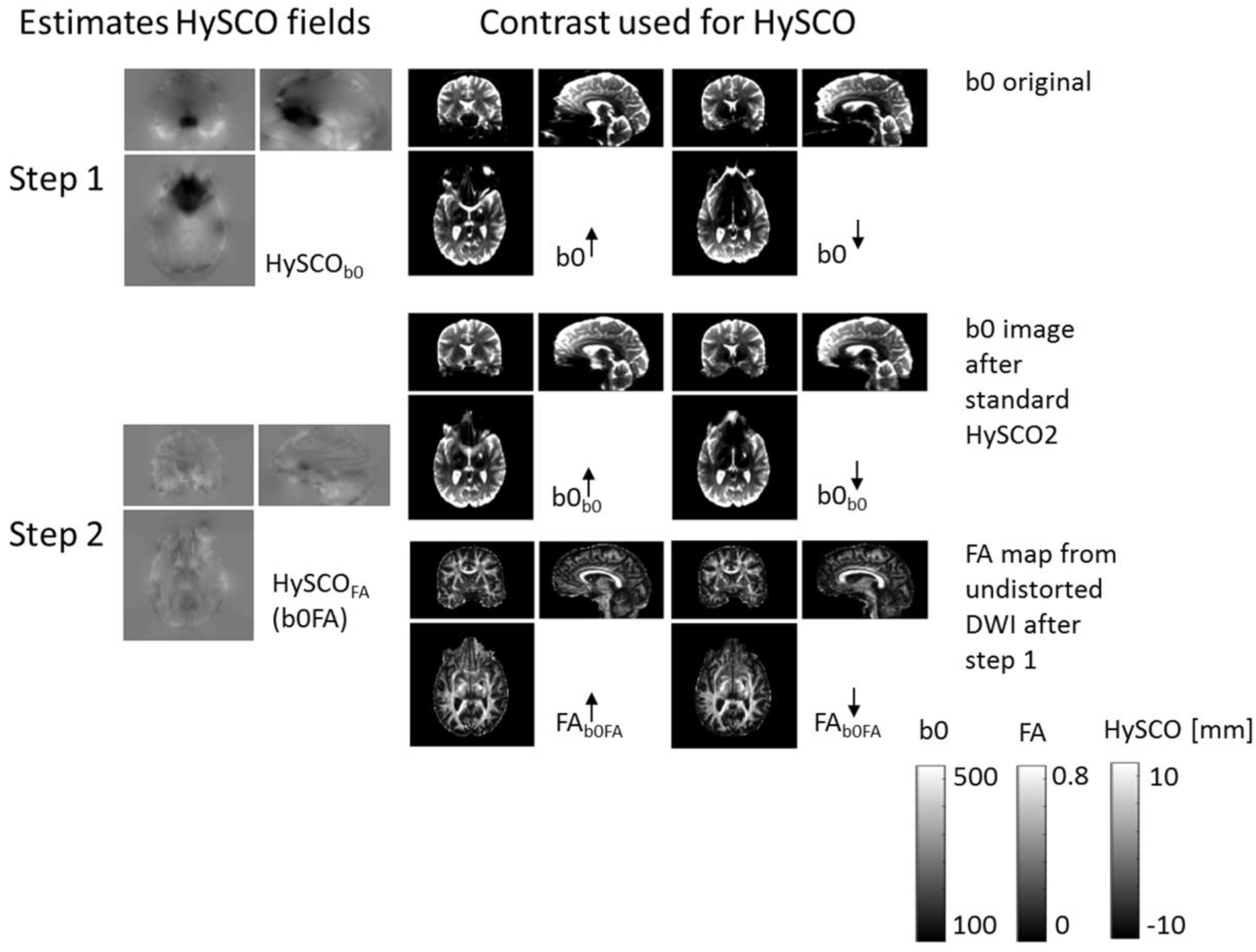
The new consecutive HySCO pipeline. In step 1, as per the standard HySCO pipeline, the HySCO module is applied to the original b0 images, estimating the HySCO field (HySCO_b0_). In step 2, tensor fitting is performed on the distortion corrected blip-up and blip-down data (b0_b0_) to estimate Fractional Anisotropy maps (FA_b0FA_), which are then used in a second round of HySCO for estimating the residual susceptibility related distortions HySCO_FA_ (b0FA). The HySCO field map is the displacement in the phase-encoding direction in mm.

#### FSL pipeline

While our main interest was in comparing the two HySCO approaches, the diffusion MRI data were also processed based on the widely-used human connectome project pipeline (Glasser et al., 2013) in FSL 6.0.1 (Smith et al., 2004) to provide external benchmarking. The blip-up and blip-down data were separately corrected for distortions using TOPUP (Andersson et al., 2003), and this was then used as the input into eddy, with the distortion, motion and eddy-current correction performed together (Andersson and Sotiropoulos, 2016) with the replace outliers option enabled (Andersson et al., 2016).

### Diffusion MRI phase-encoding combination methods

For both the consecutive HySCO and FSL methods, the distortion-corrected data sets with opposite phase-encoding polarity were combined using the ACID toolbox within SPM12.

First, the consecutive HySCO blip-up and blip-down distortion corrected data were combined by calculating the AM of each blip-up and blip-down image pair.

Second, the consecutive HySCO blip-up and blip-down distortion corrected data were combined using a WA. This aimed to minimise information loss due to susceptibility distortion blurring induced by local spatial compression. To maximise the effective spatial resolution when combining the two images acquired with opposite phase-encoding directions, a region that was stretched in one of the images, increasing effective resolution, was up-weighted whereas the opposite image was down-weighted using a WA:

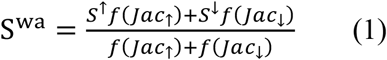

with 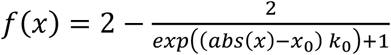 being a Fermi function with two parameters (*x*_0_, *k*_0_), tuning the sensitivity of the weighting of the Jacobian in a given voxel. Both parameters were heuristically optimized. Note that the Jacobian of the field map estimated in the initial HySCO step (Step 1, Figure 2) was used for weighting. The Jacobians were defined as follows:

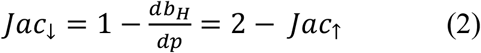

with *b*_*H*_ being the HySCO field in units of displacement in the phase-encoding direction in mm and *p* being the phase-encoding direction – here the anterior-posterior direction.

Finally, to provide a benchmark, the FSL blip-up and blip-down distortion corrected data were combined by calculating the AM of each blip-up and blip-down image pair.

### Structural MRI data pre-processing

The structural MRI data were processed for each participant using the hMRI toolbox (Tabelow et al., 2019) within SPM12. The default toolbox configuration settings were used, with the exception that correction for imperfect spoiling was additionally enabled (Corbin and Callaghan, 2021). The output MT saturation map, used as the anatomically faithful structural image, quantified the degree of saturation of the steady state signal induced by the application of the off-resonance pre-pulse, having accounted for spatially varying T_1_ times and RF field inhomogeneity (Helms et al., 2008; Weiskopf et al., 2013).

Each participant’s MT saturation map was segmented into white matter probability maps using the unified segmentation approach (Ashburner and Friston, 2005), but with no bias field correction (since the MT saturation map does not suffer from any bias field modulation) and using the tissue probability maps developed by Lorio et al. (2016).

### Comparison of the diffusion MRI distortion correction methods

Analyses to investigate the differences between the diffusion MRI distortion correction methods were performed in native space. For each participant, two FA maps were estimated for each distortion correction method (standard HySCO, consecutive HySCO, FSL); one FA map from the distortion corrected blip-up data and the other from the distortion corrected blip-down data. FA maps were estimated using the Diffusion Kurtosis Fit (DKI), utilising all the b-values, in the ACID toolbox. Each FA map was then co-registered to an individual’s MT saturation white matter probability map using a rigid-body transformation. The difference between the individual blip-up and blip-down FA maps was calculated, restricted to a white-matter mask. The white-matter mask was generated by thresholding the MT saturation white matter probability maps above 0.9. From the resulting difference image, the root mean square over all voxels in the white matter mask was calculated for each individual for each distortion correction method.

Two-tailed paired t-tests, calculated in SPSS v25 and thresholded at p < 0.05, were then performed to test for significant differences between the root mean square for each of the distortion correction methods across participants. Effect sizes are reported as Hedge’s g_av_, which is Cohen’s d calculated for repeated measures and corrected for the positive bias caused by using sample estimates, using the resources provided by Lakens (2013). Paired t-tests were used instead of a repeated measures ANOVA to treat each distortion correction method independently.

### Comparison of the diffusion MRI phase-encoding combination methods

Analyses to compare the effects of the phase-encoding combination methods were also performed in native space. For each participant, three pairs of FA and b0 maps were estimated, one pair for each phase-encoding combination method (consecutive HySCO AM, consecutive HySCO WA, FSL). The FA and b0 maps were estimated using DKI fit in the ACID toolbox.

Multichannel segmentation, using the unified segmentation approach (Ashburner and Friston, 2005), but with the tissue probability maps developed by Lorio et al. (2016), was performed on each FA and b0 map pair to define the white matter tissue segments. These diffusion-derived white matter probability maps were then co-registered to each individual’s MT saturation white matter probability map using a rigid-body transformation. This allowed for comparison between the white matter probability map defined from the distortion corrected diffusion images, and the white matter probability map from the MT saturation structural map. The percentage overlap between the diffusion white matter probability map and the structural white matter probability map was then calculated for each phase-encoding combination method.

As well as investigating the overlap of the diffusion and structural white matter probability maps as a whole, we also focused our analyses on regions that suffered from greater distortion. To identify these areas, for each participant, a map of the Jacobians was calculated from the first HySCO iteration. The co-registration transformations determined above were applied to each participant’s Jacobian map to co-register the Jacobian and MT saturation structural images. The percentage overlap of the diffusion and structural white matter probability maps was calculated in regions of higher distortions. The masks for the aforementioned regions were determined by thresholding the Jacobian map at values higher than 10%, 20% and 30% deviation from unity.

Two-tailed paired t-tests, calculated in SPSS v25 and thresholded at p < 0.05, were performed to test for significant differences between the percentage overlaps for each of the phase-encoding methods (consecutive HySCO AM, consecutive HySCO WA, FSL), at each level of distortion (whole white matter map, 10%, distortion, 20% distortion, 30% distortion). As with the distortion correction methods analyses, effect sizes are reported as Hedge’s g_av_, calculated using the resources provided by Lakens (2013), and paired t-tests were used to treat each phase-encoding method independently.

### Application to g-ratio mapping

Our final set of analyses aimed to assess any benefits of the consecutive HySCO WA approach on g-ratio mapping by comparing it to the use of consecutive HySCO AM on the generation of g-ratio maps. As neuroscientific studies utilising the g-ratio are typically performed at the group level, analyses were performed in group space. The g-ratio was calculated from axonal-water fraction (AWF) and MT saturation maps according to Ellerbrock and Mohammadi (2018):

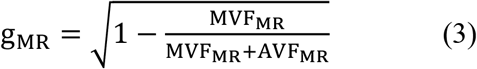

with MVF_MR_ being the myelin-volume fraction estimated from the MT saturation map and AVF_MR_ being the axonal-volume fraction. The AVF_MR_ was estimated as AVF_MR_ = (1 − MVF_MR_) AWF according to Stikov et al. (2015), where AWF was obtained by combining the intra-cellular fraction (ν_icvf_) and isotropic fraction (ν_iso_) maps from the neurite orientation dispersion and density imaging (NODDI) toolbox (http://mig.cs.ucl.ac.uk/index.php?n=Tutorial.NODDImatlab; Zhang et al., 2012) as AWF = (1 − ν_iso_) ν_icvf_. The MT saturation map was obtained from the hMRI toolbox as described above. For calibration of the MT saturation map to a myelin-volume fraction map (MVF_MR_ = α MT_sat_), we used the g-ratio based calibration method as reported in Ellerbrock and Mohammadi (2018) and Mohammadi and Callaghan (2020), with a reference g-ratio value of 0.7 in the splenium (Mohammadi et al., 2015a), yielding α = 0.217.

Transformation from native to group space was implemented using the voxel-based quantification (VBQ) approach as implemented in the hMRI toolbox (Draganski et al., 2011), which allows for preservation of the quantitative values of the g-ratio maps during smoothing (Tabelow et al., 2019). Briefly, inter-subject registration using DARTEL (Ashburner, 2007) was performed on the segmented MT saturation grey and white matter probability maps. The resulting DARTEL template and deformations were then used to normalize the MT saturation, g-ratio and Jacobian maps to Montreal Neurological Institute (MNI) space at 1.5 × 1.5 × 1.5mm resolution with a tissue weighted smoothing kernel of 6mm full width at half maximum.

Differences in g-ratio mapping were determined by comparing the calculated g-ratio values. As the g-ratio can only be defined when the white matter of the diffusion and structural (MT saturation) maps overlap, a g-ratio of zero is recorded for non-overlapping voxels (Mohammadi and Callaghan, 2020). We predicted, therefore, that the poorer overlap between the AM diffusion data and structural white matter map (due to the higher blurring in the AM diffusion data) would increase the number of voxels with a g-ratio of zero per participant, decreasing the averaged g-ratio values at the group-level in regions with high susceptibility-related distortions.

To quantify the reduction of group-averaged g-ratio values within regions suffering from high levels of distortion, for each participant we extracted the mean g-ratio from the AM and WA g-ratio maps from regions previously identified as suffering from distortion. This was performed using a mask created from thresholding the Jacobian map at 20% (see also Comparison of the diffusion MRI phase-encoding combination methods section above). We then compared these to the averaged g-ratio value in white matter using two tailed paired t-tests, thresholded at p < 0.05, calculated in SPSS v25 with effect sizes reported as Hedge’s g_av_, calculated using the resources provided by Lakens (2013). We hypothesised that smaller g-ratio values would be apparent in the AM-based g-ratio map because smoothing with zero-valued g-ratio map voxels, indicative of misalignment, would systematically lower the g-ratio values.

Finally, to localise the voxels where the AM-based g-ratio values were smaller than the WA-based g-ratios, we performed a voxel-wise comparison of the AM and WA g-ratio maps within the same 20% Jacobian distortion mask. We did this using two-tailed paired t-tests to identify significant differences in g-ratio values at statistical thresholds of p < 0.05 family wise error (FWE) with a minimum cluster size of 10 voxels.

## RESULTS

### Comparison of diffusion MRI distortion correction pipelines

The average root mean square of the difference in FA map alignment between the blip-up and blip-down data was significantly smaller for the consecutive HySCO approach in comparison to the standard HySCO approach for all participants (Figure. 3, mean difference = 0.017, t(17) = 8.65, p < 0.001, Hedge’s g_av_ = 0.91), indicating better distortion correction by the consecutive HySCO pipeline. Of note, smaller deviations (Figure 3) were observed between the blip-up and blip-down data after FSL distortion correction in comparison to the standard HySCO approach (mean difference = 0.040, t(17) = 10.38, p < 0.001, Hedge’s g_av_ = 2.49) and the consecutive HySCO approach (mean difference = 0.023, t(17) = 8.63, p < 0.001, Hedge’s g_av_ = 1.72).

**FIGURE 3.**
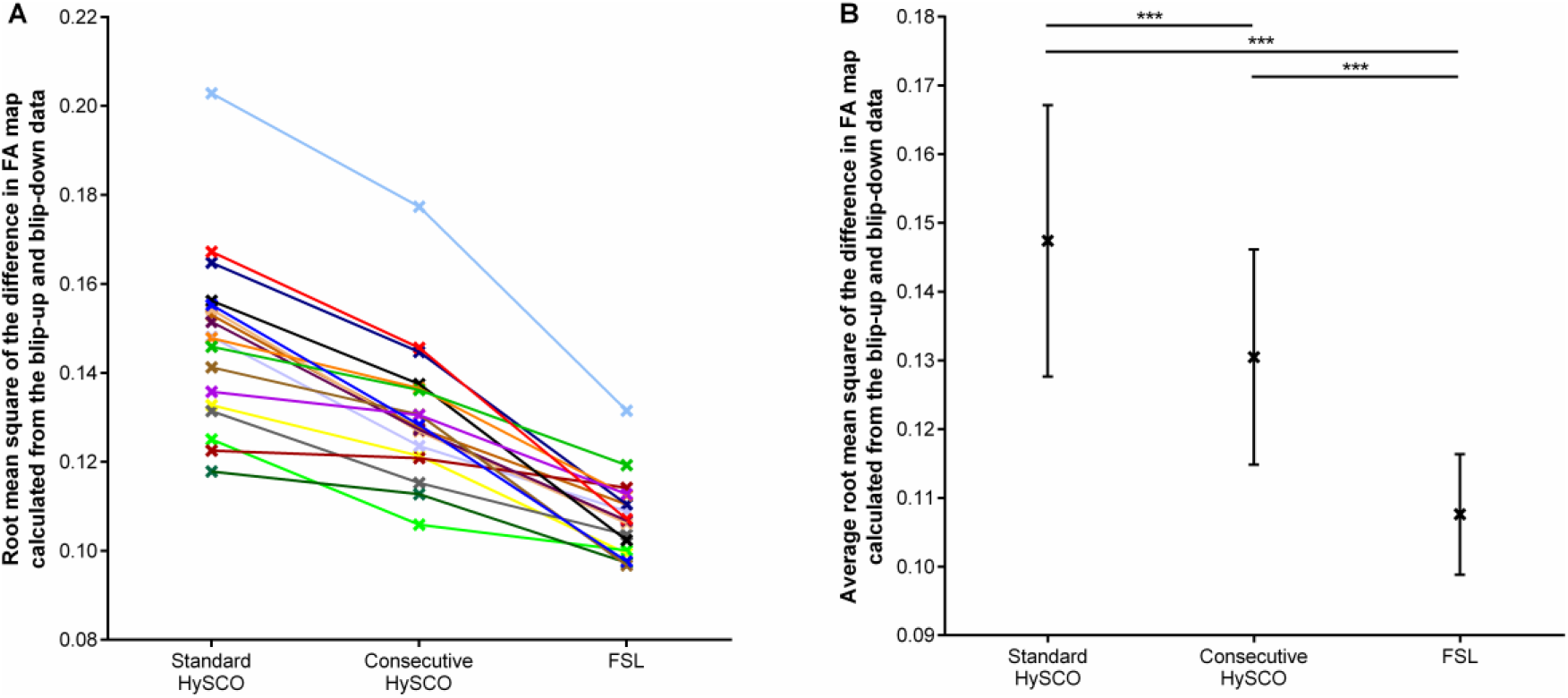
The root mean square of the difference in FA map calculated from the blip-up and blip-down data, depicted for each distortion correction methodology. **(A)** Each individual data point. **(B)** The mean and standard deviation of the sample for each distortion correction methodology. *** p < 0.001

### Comparison of diffusion MRI phase-encoding combination methods

Having established that consecutive HySCO gave better distortion correction than standard HySCO, we then turned to examining the consecutive HySCO AM and consecutive HySCO WA phase encoding combination methods. The improvement to the spatial accuracy gained by using the WA to combine the opposite phase-encoding data is shown in Figure 4. In panels A and B, additional white matter, in line with the anatomically accurate white matter tissue probability derived from the MT saturation map, can be observed in the ventromedial prefrontal cortex when using the WA compared to using the AM. In panels C and D, additional and better definition of the white matter, again in line with the anatomically accurate MT saturation white matter tissue probability map, is evident in the inferior temporal gyrus when using the WA compared to using the AM. Using the WA seems, therefore, to provide better alignment between the diffusion and quantitative structural maps.

**FIGURE 4.**
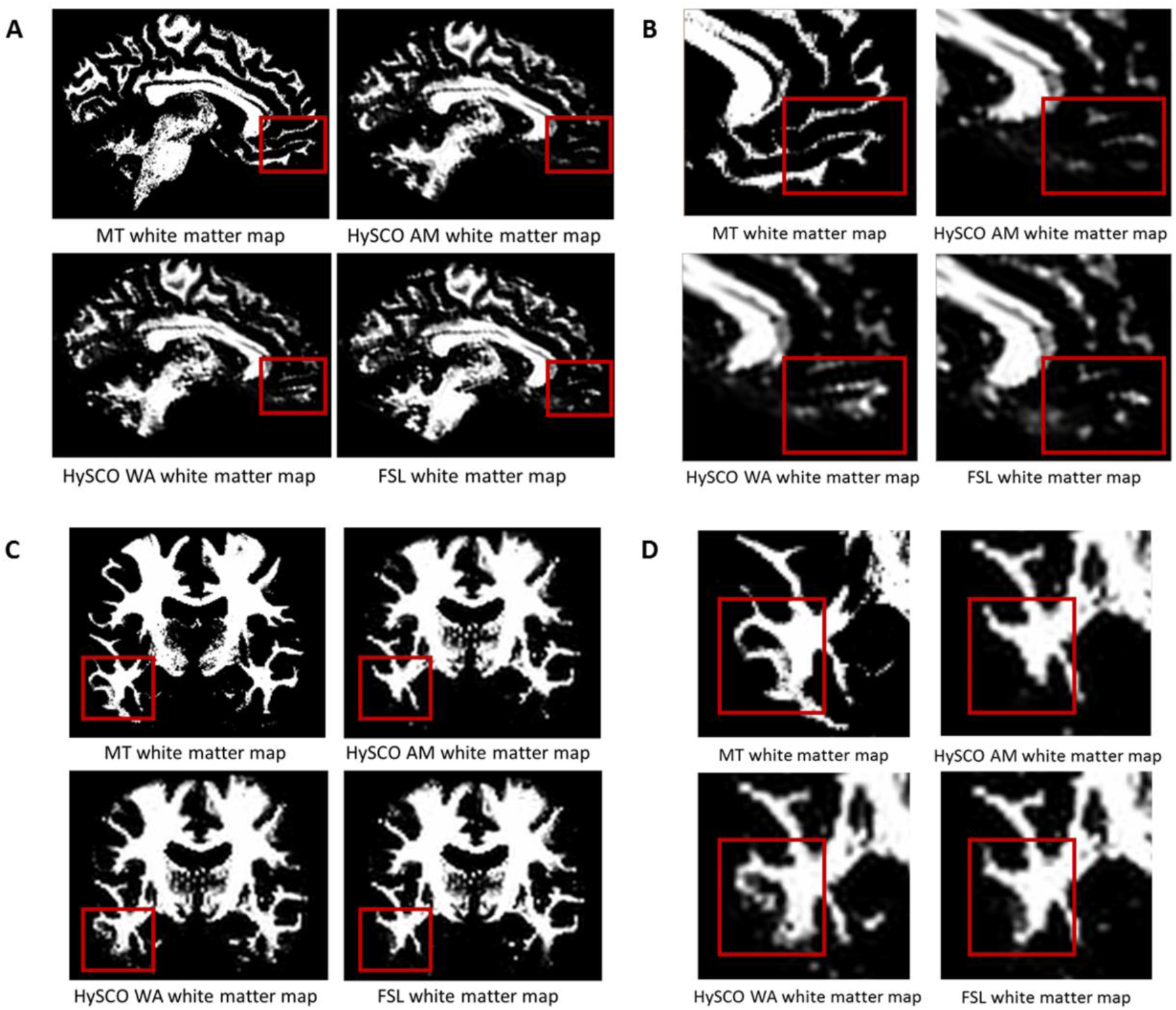
Examples of the improvements in alignment between the white matter tissue probability maps derived from the diffusion data and MT saturation map when using the WA phase-encoding combination method compared to the AM. **(A)** In one example participant (native space) additional white matter is evident in the ventromedial prefrontal cortex when using the WA. **(B)** A close-up of the white matter shown in (A). **(C)** Additional and better definition of the white matter in the inferior temporal gyrus of a different example participant (native space) when using the WA. **(D)** A close-up of the white matter shown in (C). The structural white matter map is the white matter tissue probability map derived from the MT saturation map. HySCO = consecutive HySCO, AM = arithmetic mean, WA = weighted average.

To provide a quantitative examination, we also calculated the mean percentage overlap between the diffusion and MT saturation white matter tissue probability maps for the consecutive HySCO AM, consecutive HySCO WA and, as an additional benchmark, FSL (see Figure 5). This quantitative comparison supported the qualitative visual inspection.

**FIGURE 5.**
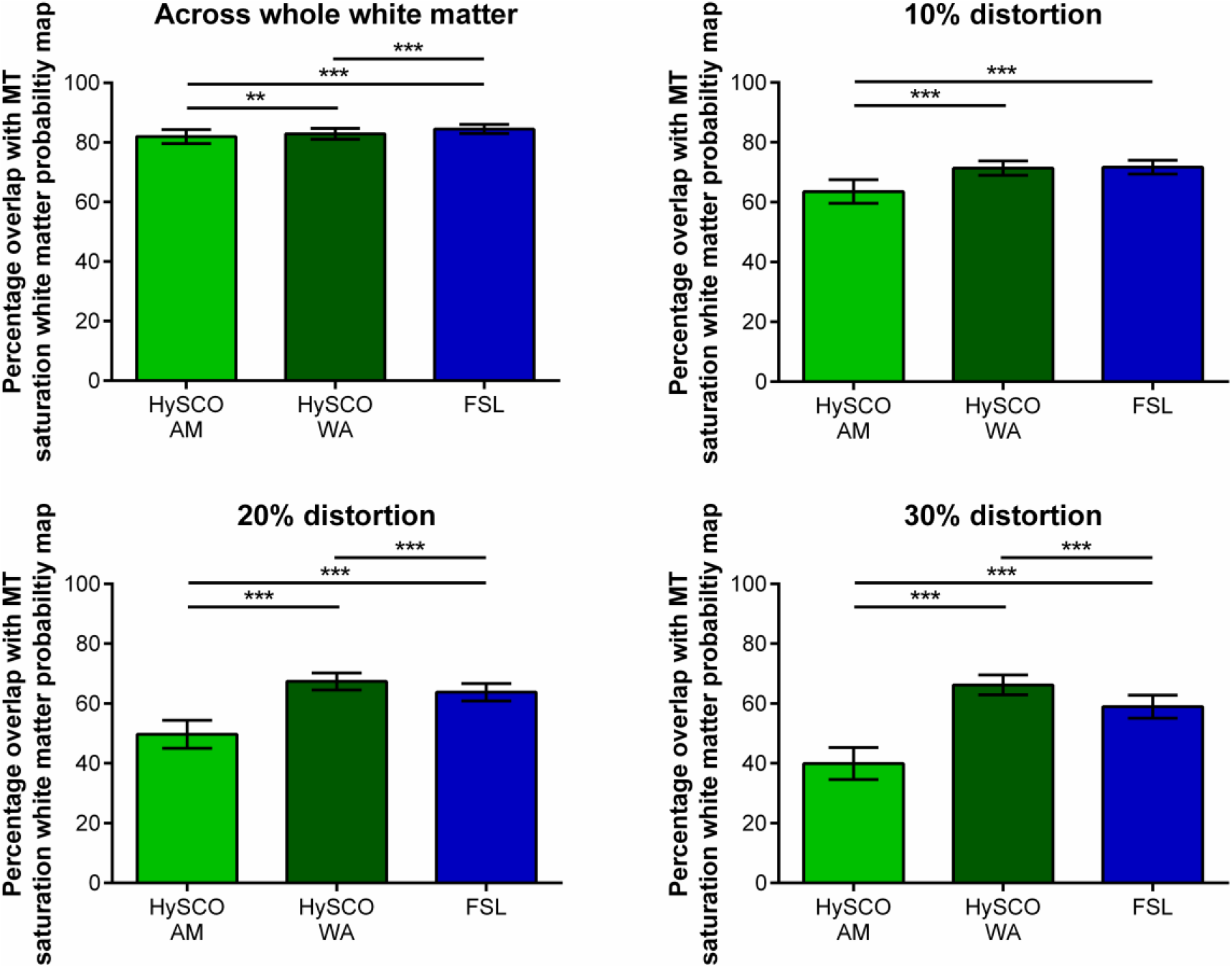
Mean (± standard deviation across participants) percentage overlap of the white matter probability maps created from the different phase-encoding combination diffusion data with the MT saturation white matter tissue probability map. The 10%, 20% and 30% distortion graphs focus on regions suffering from greater distortion, determined by thresholding the Jacobian map at distortion values higher than 10%, 20% and 30% respectively. HySCO = consecutive HySCO, AM = arithmetic mean, WA = weighted average. ** p < 0.01, *** p < 0.001.

Using a WA to combine the consecutive HySCO distortion corrected blip-up and blip-down data resulted in greater percentage overlap between the diffusion and MT saturation white matter tissue probability maps when compared to the consecutive HySCO AM. This was particularly evident in regions suffering from high levels of distortion (whole white matter: mean difference = 0.89, t(17) = 3.10, p = 0.007, Hedge’s g_av_ = 0.40; 10% distortion: mean difference = 7.81, t(17) = 11.92, p < 0.001, Hedge’s g_av_ = 2.27; 20% distortion: mean difference = 17.66, t(17) = 20.73, p < 0.001, Hedge’s g_av_ = 4.38; 30% distortion: mean difference = 26.23, t(17) = 25.94, p < 0.001, Hedge’s g_av_ = 5.64). Of note, using the AM to combine the FSL distortion corrected blip-up and blip-down data also resulted in greater percentage overlap between the diffusion and MT saturation white matter tissue probability maps compared to using the AM to combine the consecutive HySCO distortion corrected data (whole white matter: mean difference = 2.53, t(17) = 8.65, p < 0.001, Hedge’s g_av_ =1.22; 10% distortion: mean difference = 8.15, t(17) = 13.50, p < 0.001, Hedge’s g_av_ = 2.40; 20% distortion: mean difference = 14.09, t(17) = 16.88, p < 0.001, Hedge’s g_av_ = 3.46; 30% distortion: mean difference = 18.97, t(17) = 17.42, p < 0.001, Hedge’s g_av_ = 3.92).

Interestingly, across the whole of the white matter map, a larger percentage overlap between the diffusion and MT saturation white matter tissue probability maps was observed when using the AM to combine the FSL distortion corrected blip-up and blip-down data compared to the consecutive HySCO WA (mean difference = 1.64, t(17) = 6.47, p < 0.001, Hedge’s g_av_ = 0.88), although there was no difference between the methodologies at 10% distortion (mean difference = 0.34, t(17) = 0.68, p = 0.51, Hedge’s g_av_ = 0.13). However, we found that using the WA to combine the consecutive HySCO distortion corrected blip-up and blip-down data resulted in greater percentage overlap of the diffusion and MT saturation white matter tissue probability maps compared to using the AM to combine the FSL distortion corrected data in regions suffering from greater levels of distortion (20% distortion: mean difference = 3.58, t(17) = 6.13, p < 0.001, Hedge’s g_av_ = 1.19; 30% distortion: mean difference = 7.26, t(17) = 9.62, p < 0.001, Hedge’s g_av_ = 1.93) (Figure 5).

The consecutive HySCO WA method seems, therefore, to be particularly advantageous in regions suffering from high distortion, for example, in the ventromedial prefrontal cortex (as shown in Figure 4A and B), inferior temporal gyri (Figure 4C and D), as well as the middle temporal gyri, medial temporal lobe (including the anterior hippocampi), and the brain stem (Figure 6).

**FIGURE 6.**
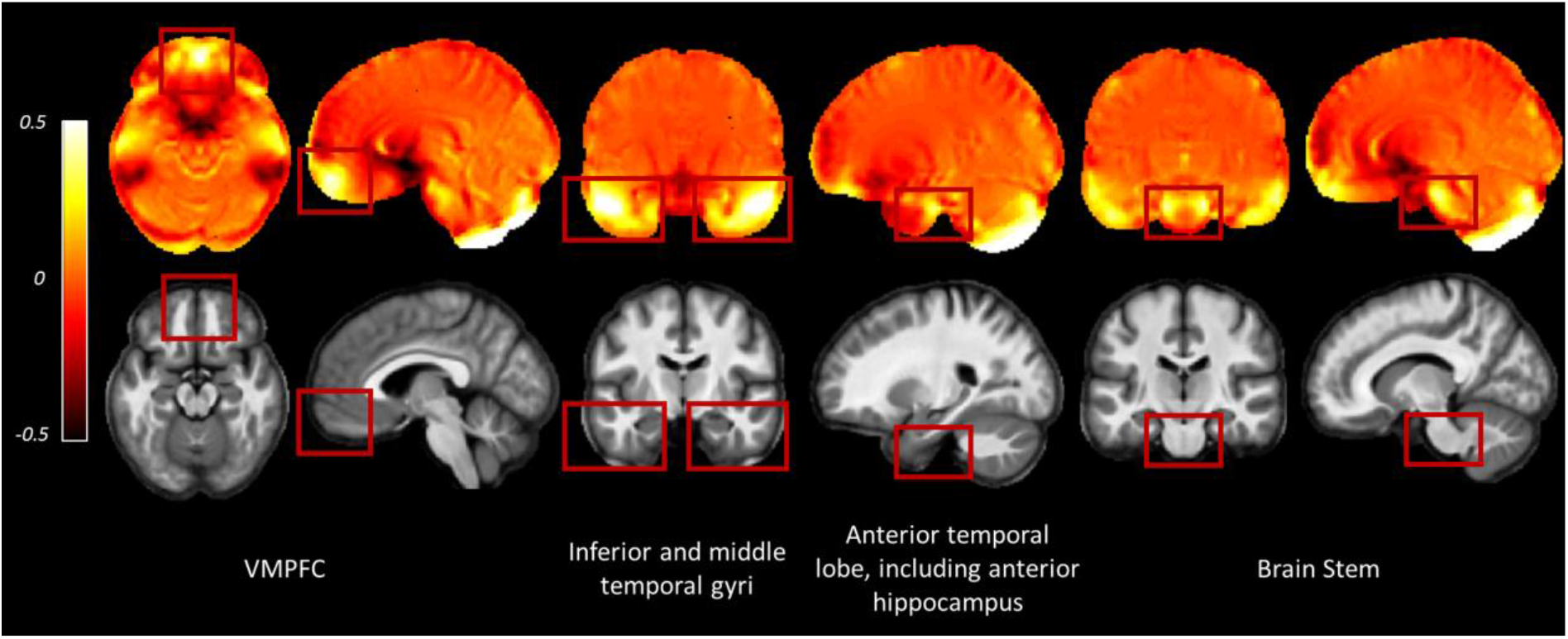
Examples of regions where greater distortion was observed across the participants. Images are the average deviation from unity of the Jacobian maps (top row) and average MT saturation map (bottom row) across participants for the whole sample in Montreal Neurological Institute (MNI) space. In the deviation from unity of the Jacobian maps, values away from zero indicate areas suffering from greater distortion (see equation 2). Inter-subject registration was performed using DARTEL as implemented in SPM12 (Ashburner, 2007). The DARTEL template and deformations were used to normalize each participant’s Jacobians and MT saturation map to the stereotactic space defined by the MNI template (at 1.5 × 1.5 × 1.5 mm resolution). VMPFC = ventromedial prefrontal cortex.

### Application to g-ratio mapping

Finally, we set out to assess whether there was any benefit of using the consecutive HySCO WA diffusion data over the consecutive HySCO AM data for g-ratio mapping. Figure 7 shows that the coverage of the g-ratio map across white matter is enhanced when using the WA combination compared to the AM combination. The focus is the ventromedial prefrontal cortex and the brain stem, two regions that suffer from high levels of distortion. As can be observed, greater accuracy was obtained in the white matter probability maps and more information was available in the AWF map. Consequently, this resulted in a larger area where the g-ratio could be defined when using the diffusion data corrected using the WA compared to performing the same calculations on the AM combination.

**FIGURE 7.**
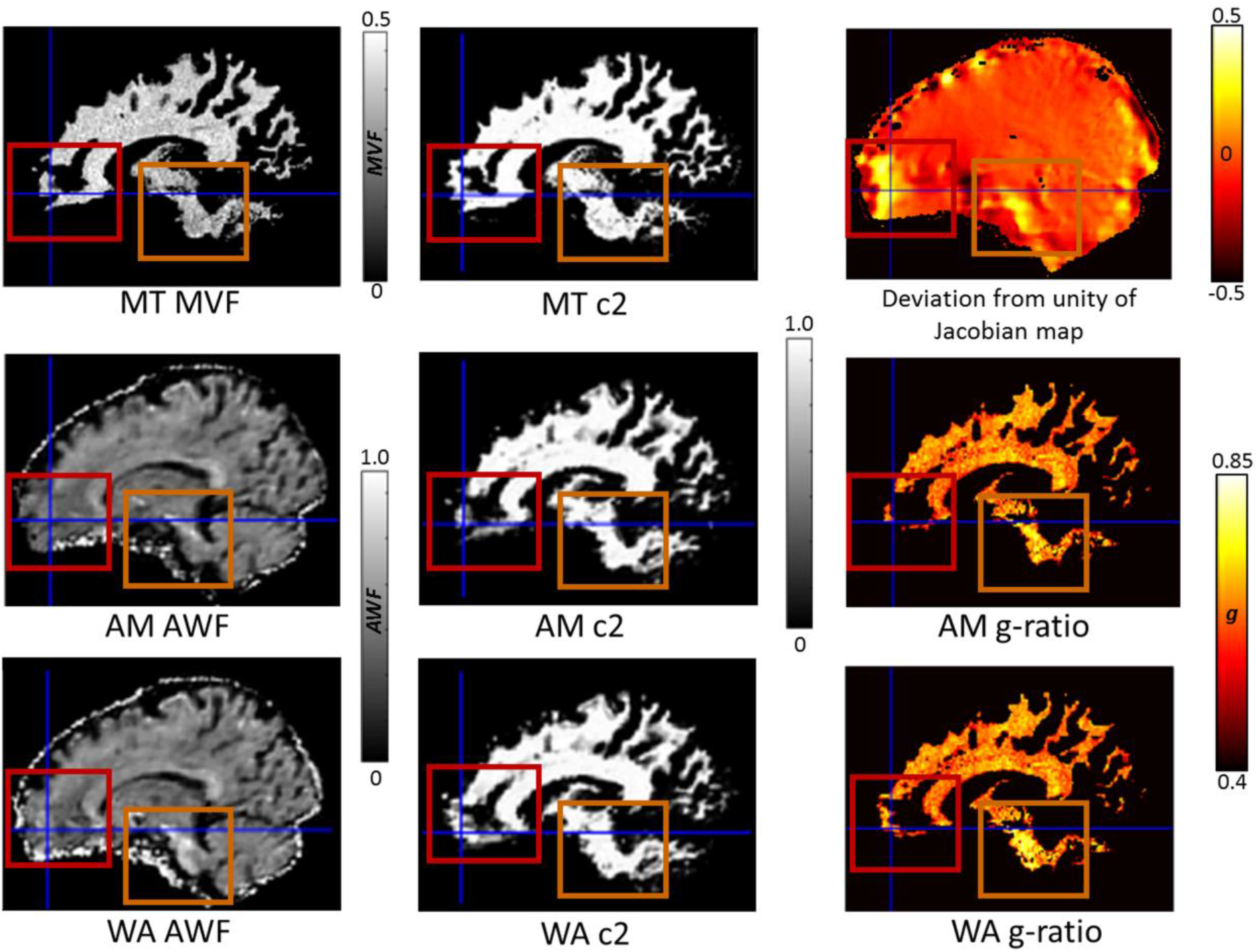
The impact of reduced blurring artefacts on g-ratio mapping when using the weighted average (WA) combination compared to the arithmetic mean (AM), illustrated for a single participant, in their native space. Depicted in the 1^st^ column are the constituents of the g-ratio: myelin-volume fraction (MVF) based on the MTsat (top), axonal-water fraction (AWF) using AM (middle), and AWF using WA (bottom). In the 2^nd^ column are the associated white matter probability maps (c2): MT c2 (top), AM c2 (middle), and WA c2 (bottom). In the 3^rd^ column is the deviation from unity of the Jacobian map (top, where values away from zero indicate areas suffering from greater distortion, see equation 2) and the resulting g-ratio maps AM g-ratio (middle), and WA g-ratio (bottom). The red boxes and crosshairs highlight the ventromedial prefrontal cortex, and the orange boxes the brain stem, both of which suffer from high levels of distortion. Greater accuracy was obtained in the white matter probability maps when using the WA phase-encoding combination method, resulting in a larger area where the g-ratio was defined.

We next sought to examine the g-ratio mapping data at the group level, predicting higher mean g-ratio values when there was better overlap between the diffusion data and the MT saturation white matter tissue probability map. As our previous analyses identified that the WA combination resulted in greater overlap between the diffusion and MT saturation white matter tissue probability maps in areas suffering from high levels of distortion, we predicted that improvements in g-ratio mapping would be evident when using the WA methodology in such regions.

To investigate this, we first compared the mean g-ratio values from across the whole of white matter, finding significantly greater WA-based g-ratios in comparison to the AM-based g-ratios (WA-based mean = 0.55; AM-based mean = 0.54; mean difference = 0.0071, t(17) = 4.58, p < 0.001, Hedge’s g_av_ = 0.45). To create a non-biased baseline mean g-ratio value, for each participant we calculated a combined mean g-ratio for the AM and WA white matter maps. We then extracted the mean g-ratio from the regions suffering from high levels of distortion (identified by thresholding the Jacobian map at 20%), and subtracted this from the combined mean g-ratio in order to provide a measure of the difference in g-ratio between highly distorted regions and the whole of white matter. We predicted that the difference in mean g-ratio values between the highly distorted regions and the combined mean g-ratio across the whole of the AM and WA white matter maps would be smaller for the WA-based g-ratios. This is because we expected the g-ratio values in the AM-based g-ratio map to be lower in highly distorted regions, and thus further away from the combined mean g-ratio. This would be due to the poorer overlap between the diffusion and MT saturation white matter tissue probability maps increasing the number of voxels with a g-ratio of zero.

Indeed, a smaller difference between the g-ratio values in highly distorted regions and the combined mean g-ratio was apparent for the WA-based g-ratios compared to the AM-based g-ratios (mean difference = 0.092, t(17) = 12.76, p < 0.001, Hedge’s g_av_ = 1.98; Figure 8A). This suggests, therefore, that in regions suffering from high levels of distortion, the WA-based g-ratio values were closer to the white-matter average than AM-based g-ratio values, thus providing a better representation of g-ratio values in regions with high susceptibility-related distortions.

**FIGURE 8.**
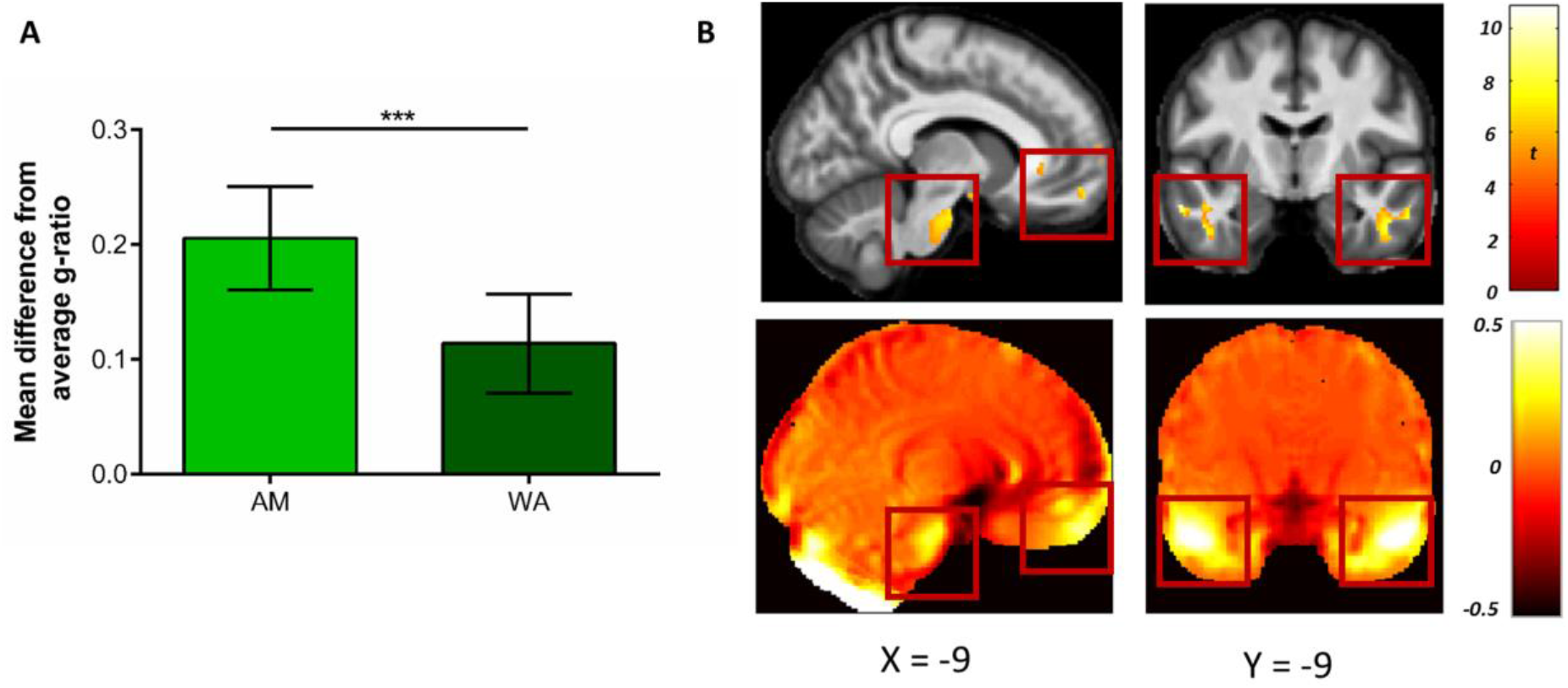
Comparison of g-ratio values when using the weighted average (WA) and arithmetic mean (AM) combinations. **(A)** Mean (± standard deviation) difference in g-ratios between highly distorted regions (defined by the 20% Jacobian distortion mask) and the whole of white matter. The smaller difference evident after the WA combination suggests the g-ratio values were closer to the white matter average, providing a better representation of g-ratio values in regions with high susceptibility-related distortions. **(B)** Voxel-wise comparison within the 20% Jacobian distortion mask showing regions with significantly higher g-ratio values when using the WA in comparison to the AM combination (top row), and their overlap with regions suffering from high levels of distortion. This is observable via the deviation from unity of the Jacobian map (bottom row, where values away from zero indicate areas suffering from greater distortion, see equation 2). Comparison images are thresholded at p < 0.05 FWE corrected and displayed on the average MT saturation maps for the whole sample. The left images show the ventromedial prefrontal cortex and brain stem, the right images show the inferior and middle temporal gyri.

Finally, to further localise where, within the regions with high susceptibility-related distortions, the AM-based g-ratios were significantly smaller than the WA-based g-ratios, voxel-wise statistics were performed. As can be seen in Figure 8B, higher g-ratio values were identified (top row) when using the WA combination in regions suffering from the greatest levels of distortion (bottom row), including the ventromedial prefrontal medial frontal cortex, brain stem, and inferior and middle temporal gyri (see also Table 1). No significant voxels were identified for the reverse contrast.

**TABLE 1.**
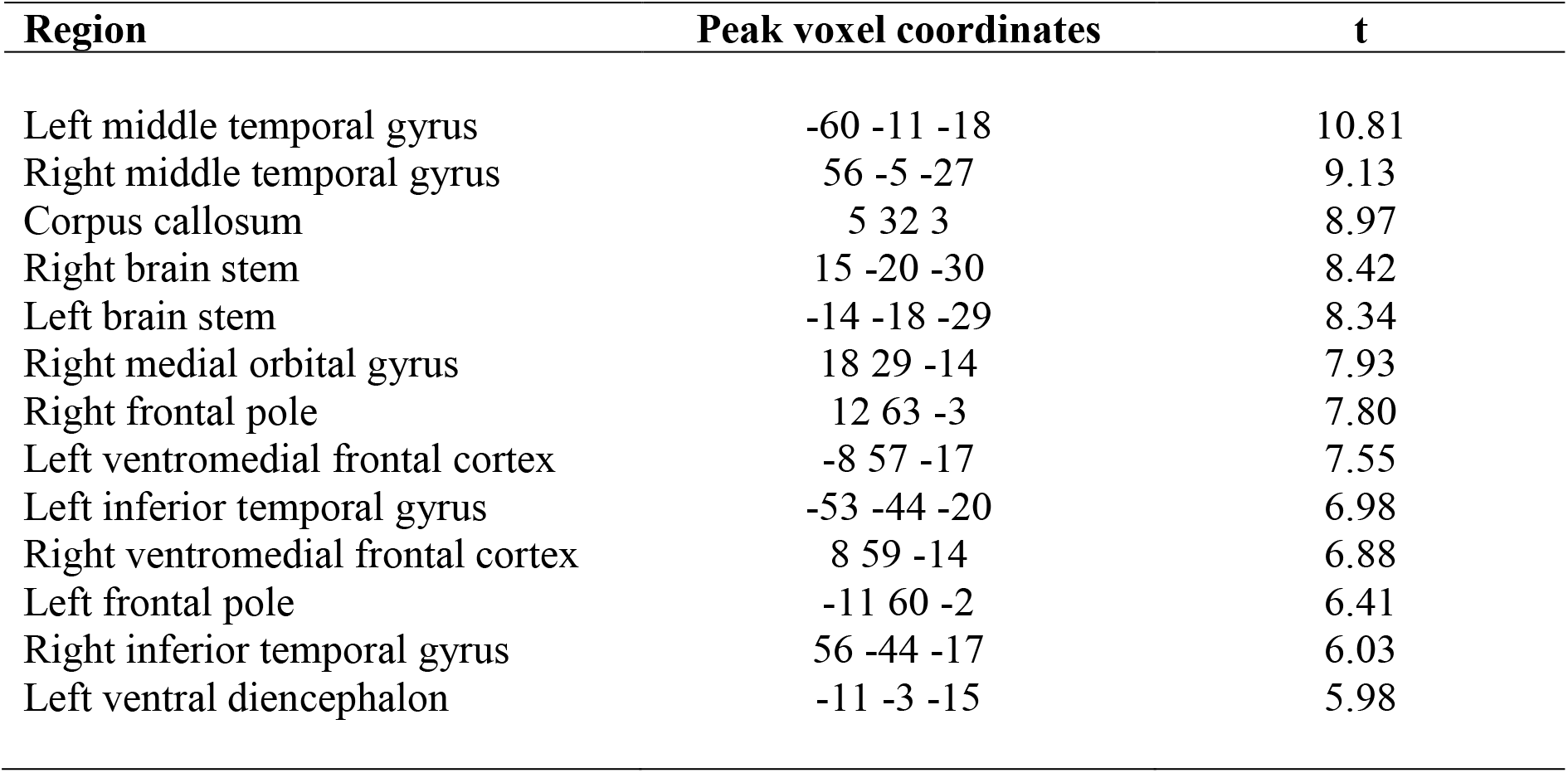
Regions and peak voxel coordinates with significantly (p < 0.05 FWE corrected) higher g-ratio values within the 20% distortion mask when using the weighted average combination compared to the arithmetic mean.

| Region | Peak voxel coordinates | t |
| --- | --- | --- |
| Left middle temporal gyrus | -60 -11 -18 | 10.81 |
| Right middle temporal gyrus | 56 -5 -27 | 9.13 |
| Corpus callosum | 5 32 3 | 8.97 |
| Right brain stem | 15 -20 -30 | 8.42 |
| Left brain stem | -14 -18 -29 | 8.34 |
| Right medial orbital gyrus | 18 29 -14 | 7.93 |
| Right frontal pole | 12 63 -3 | 7.80 |
| Left ventromedial frontal cortex | -8 57 -17 | 7.55 |
| Left inferior temporal gyrus | -53 -44 -20 | 6.98 |
| Right ventromedial frontal cortex | 8 59 -14 | 6.88 |
| Left frontal pole | -11 60 -2 | 6.41 |
| Right inferior temporal gyrus | 56 -44 -17 | 6.03 |
| Left ventral diencephalon | -11 -3 -15 | 5.98 |

## DISCUSSION

Our aims in this study were to improve the susceptibility-related distortion correction for diffusion MRI data in SPM, to deploy an alternative approach to reducing image blurring in diffusion MRI data when combining blip-up and blip-down EPI data after susceptibility-related distortion correction, and to assess any benefits of these changes for g-ratio mapping. We found that the new processing pipeline, consecutive HySCO, improved distortion correction. Moreover, the WA compared to the commonly used AM combination method showed a distinct benefit following distortion correction in SPM, and this performed well when benchmarked against FSL, particularly in brain areas with strong susceptibility-related distortions. Finally, we showed that the consecutive HySCO WA performed better than the AM method when combined with other quantitative MRI parameters – MPM – to perform g-ratio mapping. These improvements mean that researchers can conveniently access a wide range of diffusion-related analysis methods within one framework because they are now available within the open-source ACID toolbox as part of SPM, which can be easily combined with other SPM toolboxes, such as the hMRI toolbox, to facilitate computation of myelin biomarkers that are necessary for g-ratio mapping.

### The new diffusion MRI distortion correction pipeline in ACID

Recent work has highlighted that while the distortion correction module in SPM-ACID (HySCO2) performed well in correcting distortion in b0 images, there were limitations in its ability to reduce susceptibility-related distortion (Gu and Eklund, 2019), highlighting a need for improvement. To enhance the performance of the HySCO2 module, we performed two consecutive applications of the HySCO2 module instead of the standard single application, with ECMOCO for motion and eddy current correction performed in both the standard and consecutive HySCO pipelines. The second iteration of HySCO2 in the consecutive HySCO pipeline used FA maps computed after the initial distortion correction as input. We found that the consecutive HySCO pipeline improved distortion correction as compared to the standard HySCO pipeline, while noting that it was less efficient than the FSL TOPUP and eddy pipeline.

### WA combination – a method to reduce susceptibility-distortion related blurring artefacts

The new consecutive HySCO pipeline was used to test the performance of an alternative phase-encoding combination approach. Areas that are strongly squeezed by susceptibility-related distortions suffer from blurring artefacts when taking the AM of the susceptibility-related distortion corrected blip-up and blip-down images. We proposed the WA combination to reduce this blurring artefact. The WA combination uses the Jacobian of the distortion field (estimated by the HySCO2 module in ACID during susceptibility-related distortion correction) to down-weight areas that are squeezed in, for example, the blip-up image, while up-weighting the same (stretched) regions in the opposite blip-down image. As illustrated in Figure 4, compared to the consecutive HySCO AM, using the consecutive HySCO WA combination resulted in the identification of additional and better spatially defined white matter, in line with the more anatomically accurate MT saturation white matter tissue probability map that was used here as a reference.

The benefits of the WA phase-encoding combination method were also apparent when using quantitative comparisons. Comparing the use of the WA and AM combination methodologies (following distortion, eddy current and motion correction in SPM) identified an increase in the percentage overlap of the diffusion and MT saturation white matter tissue probability maps when using the WA. This was apparent across the whole white matter map, but particularly when focusing on regions suffering from high levels of distortion (e.g. ventromedial pre-frontal cortex, inferior and middle temporal gyri, medial temporal lobe, brain stem). The WA also compared well with FSL, which was used for benchmarking, and indeed showed a greater percentage overlap of the diffusion and MT saturation white matter tissue probability maps for regions suffering from high levels of distortion.

### Application to g-ratio mapping

The beneficial effect of using the consecutive HySCO WA combined diffusion data was illustrated for g-ratio mapping, where blurring effects can reduce the overlap between the diffusion-MRI-based axonal biomarker and the myelin marker maps. The blurring artefacts in diffusion MRI (which are decreased when using a WA combination) lead to a worse delimitation between grey and white matter, which in turn can propagate towards the resulting g-ratio map (Mohammadi and Callaghan, 2020). As the MR g-ratio is defined only in white matter (Stikov et al., 2015), it is typically created by combining an axonal water fraction estimated in the white matter probability maps from diffusion MRI (here, the AWF) and a white matter myelin biomarker map (here, the MT saturation). In current implementations, whenever one of the two constituents is not defined, the g-ratio value cannot be calculated, resulting in undefined areas for the g-ratio map which are set to zero (see Figure 7 in Mohammadi and Callaghan, 2020). By using the consecutive HySCO WA method to combine the diffusion data, higher quality data are available to calculate the AWF, enabling the g-ratio map to cover a larger area of the white matter than when using the consecutive HySCO AM method (illustrated in our Figure 7).

As these undefined areas vary between individuals, we additionally assessed the beneficial effects of the WA over the AM combination for g-ratio mapping at the group level. We predicted that less overlap between the diffusion and MT saturation white matter tissue probability maps would increase the number of voxels with a g-ratio of zero, decreasing the local g-ratio values in the group average. Indeed, this was what we found. In regions suffering from high levels of distortion, the WA-based g-ratio values were closer to the white-matter average than the AM-based g-ratio values. This prediction was further supported by the observation that the AM-based g-ratio values were significantly smaller than WA-based g-ratio values in regions with high susceptibility-related distortions. The improved definition of white matter in regions suffering from high levels of distortion by the WA technique seems, therefore, to also result in better representation of g-ratio values in regions with high susceptibility-related distortions.

### Limitations and future directions

We developed and tested our consecutive HySCO WA methodology within the SPM framework to facilitate the combination of diffusion and quantitative MT saturation maps, the latter of which has been developed uniquely within SPM (Tabelow et al., 2019). As such, we aimed to improve the tools available in ACID and SPM. It may be that applying the WA methodology developed here to other susceptibility-related distortion techniques, such as FSL, would also provide further benefits, however, examining this was beyond the scope of the current study.

We note that, even with the improvements to the initial SPM distortion correction pipeline, FSL TOPUP and eddy provided better initial distortion, motion and eddy correction than the SPM equivalents. This is potentially due to the consecutive HySCO pipeline using multiple interpolation steps, introducing additional blurring effects. Future work will aim to develop a methodology to combine these interpolation steps, further increasing the effectiveness of HySCO distortion correction. Of note, however, despite the poorer performance of HySCO susceptibility-related distortion correction as compared to FSL, the WA combination in ACID overcame these initial differences in distortion correction between the two toolboxes. Further development of the distortion correction methods in HySCO may result in even greater anatomical accuracy in the future.

The consecutive HySCO WA distortion correction technique may also have implications beyond the demonstrated improvements in g-ratio mapping. For example, it could assist with improved alignment of the diffusion to anatomical data when relating tractography-based macroscopic brain connections to their cortical origin (e.g. Jbabdi et al. 2015), or for multivariate analyses such as those deployed by Draganski et al. (2011). The tools developed here are freely available in the ACID toolbox for SPM for use in other methodological developments such as these.

## Conclusions

We present a new WA approach to combining blip-up and blip-down EPI images (after susceptibility-related distortion correction) that reduces image blurring in diffusion MRI data. In addition, use of the WA method in combination with other quantitative structural MRI maps that can be processed within SPM improved g-ratio mapping at the group level in regions with high susceptibility-related distortions. This WA approach is now available in the open-source SPM-ACID toolbox together with a new processing pipeline for improved susceptibly-related distortion correction.

## DATA AVAILABILITY

On publication the tools developed as part of this article will be made freely available to download from (www.diffusiontools.com). The raw diffusion and structural MRI data are part of a larger project and will be made available once the construction of a dedicated data-sharing portal has been finalised. In the meantime, requests for the raw data can be sent to.

## ETHICS STATEMENT

This study involving human participants was approved by the University College London Research Ethics Committee (project ID: 6743/001). All participants gave written informed consent.

## AUTHOR CONTRIBUTIONS

**IC:** Methodology, Investigation, Formal analysis, Writing - original draft, Writing - review & editing. **MC:** Formal analysis, Writing - review & editing. **NW:** Resources, Writing - review & editing. **EM:** Conceptualization, Methodology, Formal analysis, Writing - original draft, Writing - review & editing, Supervision, Funding acquisition. **SM:** Conceptualization, Methodology, Formal analysis, Writing - original draft, Writing - review & editing.

## ACKNOWLEDGEMENTS

Thanks to Anna Monk, Victoria Hotchin and Gloria Pizzamiglio for assistance with data collection. Thanks also to Lars Rutthotto for helpful discussions. This manuscript has been released as a pre-print at Biorxiv (Clark et al., 2021).

